# A comparison of wild and captive Comal Springs riffle beetle (*Heterelmis comalensis*) microbiomes

**DOI:** 10.1101/2021.07.12.452104

**Authors:** Zachary Mays, Amelia Hunter, Lindsay Glass Campbell, Camila Carlos-Shanley

**Author notes:** Correspondent author: Camila Carlos-Shanley.

## Abstract

The gut microbiome is affected by host intrinsic factors, diet, environment, and strongly linked to host’s health. Although fluctuations of microbiome composition are normal, some are due to changes in host environmental conditions. When species are moved into captive environments for conservation, education, or rehabilitation, these new conditions can influence a change in gut microbiome composition. Here, we compared the microbiomes of wild and captive Comal Springs riffle beetles (*Heterelmis comalensis*) by using amplicon sequencing of the 16S rRNA gene. We found that the microbiome of captive beetles was more diverse than wild beetle microbiomes. We identified 24 Amplicon Sequence Variants (ASVs) with relative abundances significantly different between the wild and captive beetles. Many of the ASVs overrepresented in captive beetle microbiomes belong to taxa linked to nitrogen-rich environments. This is one of the first studies comparing the effects of captivity on the microbiome of an endangered insect species. Our findings provide valuable information for future applications in the management of captive populations of *H. comalensis*.

## Introduction

The wide diversity of bacteria that colonize niches in animal microbiomes can have both positive or negative impacts on host health (Cong & Zhang, 2018). Members of the microbiomes can also provide information about their microenvironment and their hosts’ external environment. For example, the presence of known methanotrophs indicate that a level of methane in an environment has reached a threshold to support them (Kip *et al*., 2010) or high abundance of bacteria with roles in the nitrogen cycle could indicate high amounts of environmental nitrogen compounds (Fonseca *et al*., 2018). However, sequencing the genomes of bacteria and searching for genes linked to metabolism is a preferred method to identify a microbe’s capabilities and can offer evidence toward an explanation for their presence.

Insect microbiomes are particularly important for helping their hosts survive in their niche environments and are often adapted to their hosts’ needs (Degli Esposti & Martinez Romero, 2017). Many insects rely on their microbiome to facilitate feeding on recalcitrant foods, provide immunity and protection against pathogens and predators, mediate intra- and interspecific communication between different species, control mating and reproductive success, and supply essential amino acids and other metabolic compounds that are absent in their diet (Engel & Moran, 2013, Guyomar *et al*., 2018, Gupta & Nair, 2020, Jing *et al*., 2020). Some insects, like mosquitoes, require bacteria to develop and will perish as larvae if they are not able to acquire the bacteria needed to signal the next stage of growth (Coon *et al*., 2014, Coon *et al*., 2017).

The Comal Springs riffle beetle (*Heterelmis comalensis*; Coleoptera: Elmidae) is a flightless, spring obligate, aquatic beetle endemic to the Edwards (Balcones Fault Zone) Aquifer spring-fed Comal and San Marcos Springs in Central Texas, USA (Brown, 1987, USFWS, 2006). This aquifer has experienced decreased flow rates during droughts, overconsumption of groundwater, and decreases in water quality, likely due to urbanization of the surrounding area (USFWS, 1997). *H. comalensis* was listed as an endangered species in 1997 because of these threats (USFWS, 1997). Since 2000, a captive assurance colony of Comal Springs riffle beetles has been housed at the San Marcos Aquatic Resources Center (SMARC), a facility within the Fish and Aquatic Conservation branch of the United States Fish and Wildlife Service.

*H. comalensis* consume biofilms grown on allochthonous terrestrial coarse particulate organic matter. These biofilms derive > 80% of their essential amino acids from bacteria (Nair *et al*., 2021). Since bacteria play a significant role in their diet, we believe that their gut microbiome has an important function in their health and development. The microbiomes of arthropods are influenced by many factors including environment, diet, and metamorphic characteristics (Engel & Moran, 2013, Guyomar *et al*., 2018, Gupta & Nair, 2020, Jing *et al*., 2020). Our research compares the microbiomes of wild and captive-held adult *H. comalensis* by 16S rRNA gene amplicon sequencing. This study will inform applied research of coleopteran microbiomes, management techniques, husbandry, and provide a baseline for *H. comalensis* microbiome composition. Given increasing evidence linking microbiomes to overall health of the host (Jin Song *et al*., 2019), our findings will be of use for developing ideal conditions for endangered invertebrates in captivity.

## Materials and Methods

### Sample collection

Wild beetle samples were hand picked off submerged wood over spring openings in Comal Springs in New Braunfels, Texas. Collections occurred on February 28, 2019 (14 beetles) and May 30, 2019 (13 beetles). Captive beetles had been collected from the wild and maintained in the refugium for five to six months prior to their randomized selection as samples for the study on May 28, 2019 (6 beetles) and November 6, 2019 (12 Beetles). SMARC staff humanely euthanized and preserved wild and captive beetles in 95% ethanol before transferring them for microbiome sequencing at the lab at Texas State University.

The SMARC captive riffle beetle environment consists of a series of opaque plastic containers customized with PVC fittings to provide fresh flowing, non-recirculated Edwards (Balcones Fault Zone) Aquifer well water between 21-23 °C and filled with a layer of conditioned leaves, rocks, and wood as biofilm food sources (Huston & Gibson, 2015). Leaves, rocks, and wood are collected from the riparian area directly above or near *H. comalensi*s habitat. Potential aquatic nuisance species are removed by hand and then materials are placed in a drying oven at 60 °C for 24 hours. Materials are then conditioned by submersion in flow-through freshwater aquaculture systems for several weeks until biofilms develop. Once degradation of the organic material is apparent, they are utilized as food sources for captive *H. comalensis*. Two wells are used to pump freshwater to the captive aquatic animals at the SMARC. These wells pull water from the artesian zone of the Edwards (Balcones Fault Zone) Aquifer near the freshwater/saline-water interface. Well LR-67-09-105 (W100-899, Hunter Road, drilled in 1970, 330 feet depth) is 0.48 miles away from the SMARC and well LR-67-09-106 (W100-900, McCarty Lane, drilled in 1970, 402 feet depth) is 1.02 miles away from the SMARC. The wells likely receive the same regional discharge that feeds Comal and San Marcos Springs with a minor influence of local recharge sources (Musgrove, 2012, Johnson *et al*., 2019).

### DNA extraction and 16S rRNA gene libraries

Out of the 45 beetles that were sent to the lab, six were used for finding a satisfactory way to extract the DNA from each beetle. The beetle microbiomes were extracted by crushing individual whole beetles with a micropestle in 450 µL of MBL solution from the QIAamp BiOstic Bacteremia DNA kit by Qiagen© (one beetle per tube). The contents were vortexed and placed on a heat block for 15 minutes at 70 °C before being transferred to a Powerbead tube and homogenized for 10 minutes in two-minute intervals. The DNA extraction followed the QIAamp BiOstic Bacteremia DNA kit protocol. The DNA concentrations were measured with a Qubit 4 fluorometer. We amplified the V4 region of the 16S rRNA gene with PCR using a set of tagged MiSeq primers [primers: forward -GTGCCAGCMGCCGCGGTAA; reverse - GGACTACHVGGGTWTCTAAT]. Each sample ran with the following conditions: initial denaturation at 95 °C for three minutes, 25 cycles of 95 °C for 30 seconds, 55 °C for 30 seconds, 72 °C for 30 seconds, and a final elongation step at 72 °C for five minutes. We cleaned the PCR products with ExoSAP-IT™ PCR Product Cleanup Reagent. We sequenced the amplicon libraries and non-template controls using the paired-end v3 chemistry in the Illumina MiSeq platform at the Department of Biology, Texas State University.

### Sequencing analysis

We eliminated sequences from samples with quality scores below 30 from the targeted sequencing results using the *DADA2 v1*.*14*.*1* package (https://benjjneb.github.io/dada2/) pipeline resulting in 33 of 45 collected beetle microbiomes. We truncated the ends of sequences, checked error rates, and removed chimeras. After a cluster analysis of 269,250 sequences, we found a total of 250 amplicon sequence variants (ASVs) and assigned taxonomy to them using the *silva_nr_v138_train_set*.*fa* training set. No ASVs were found in the non-template controls.

Statistical analyses were done using various tools in the *MicrobiomeAnalyst* web platform (Dhariwal *et al*., 2017, Chong *et al*., 2020). We excluded one captive beetle microbiome dataset with less than 1000 reads resulting in 32 microbiomes (21 wild, 11 captive) with between 1036 and 14273 reads. We filtered ASVs with counts less than four in fewer than 10% of the microbiomes to remove possible sources of contamination or sequencing error. We also normalized the dataset with the Total Sum Scaling method to account for the differences in sequencing depth among the microbiomes. For each analysis in *MicrobiomeAnalyst* we used the default settings for filtering metrics and changed only graphical settings.

We used the Linear Discriminant Analysis Effect Size (LEfSE) tool to identify ASVs with relative abundance significantly different between captive and wild beetles. The non-metric multidimensional scaling was performed in the *MicrobiomeAnalyst* software with Bray-Curtis dissimilarity. We used three different diversity estimates, Simpson, Shannon, and Choa1 to estimate alpha-diversity. The Shannon index is strongly influenced by richness and the Simpson index gives more weight to common taxa and evenness in their estimations of diversity. The Chao index considers rare taxa might be not present in the dataset. Since each diversity estimate has strengths, we decided to use multiple metrics to provide more insight into the true alpha-diversity of the microbiomes.

We used the nBLAST tool in the Integrated Microbial Genomics (IMG) online toolset to search for matches between significant ASVs and isolates that we sent to the Joint Genome Institute for whole genome sequencing. For each match, we searched through several genomes with similar average nucleotide identities to compare and speculate on the origin or purpose of the bacterial isolate from the beetles. Using a FASTA file of the ASV sequences, we searched the Ribosomal Database Project (RDP), SILVA, and the National Center of Biotechnology Information (NCBI) databases for similar 16S rRNA gene sequences. The 16S rRNA gene amplicon sequencing datasets generated in this study are available in the NCBI SRA repository under the accession numbers: PRJNA725595

## Results

At the phylum level the *H. comalensis* microbiome was dominated by Bacteroidota, Firmicutes and Proteobacteria for both groups (**Figure 1a**). *Acidobacteria* phyla was enriched in captive beetles (average relative abundance 17%) when compared to wild beetles (average relative abundance 0.6%) (**Figure 1a**). The average relative abundance of *Nitrospira* phyla was 0.01% in wild beetles and 3.36% in captive beetles. The average relative abundance of the genus *Dysgonomonas* was higher in wild beetles (average relative abundance = 27.67%) than in captive beetles (average relative abundance = 14.41%). The average relative abundance of the genus *Staphylococcus* was 12.05% and present in 76.8% of captive beetles, whereas it had only an average relative abundance of 1.73% and was present in only 9.5% of wild beetles. (**Figure 1b**).

**Figure 1:**
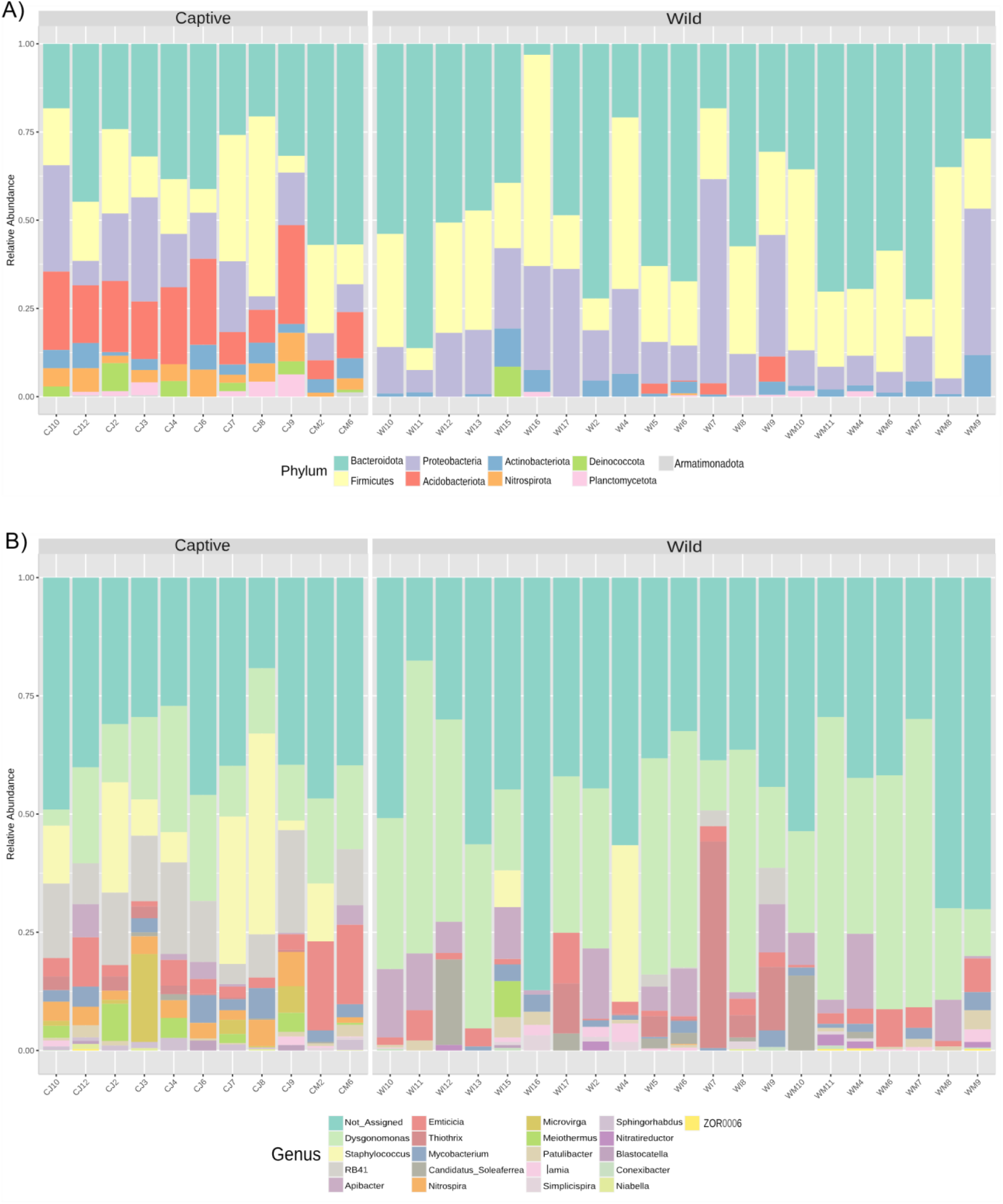
**Relative abundance bar plots** based on abundance of phyla (A) and genera (B) of the captive and wild beetle microbiomes. Only major components (abundance > 10) were included in the genera abundance bar plot.

The alpha diversity indices Chao1, Shannon, and Simpson indicated that the captive microbiomes were more diverse than the wild microbiomes on average (**Figure 2**). The differences in composition of wild and captive beetle microbiomes differed as supported by non-metric multidimensional scaling (nMDS) (PERMANOVA: F-statistic = 11.379, R^2^ = 0.27499, p-value < 0.001) (**Figure 3**). However, wild microbiomes had greater difference between groups than captive microbiomes (Figure 3).

**Figure 2:**
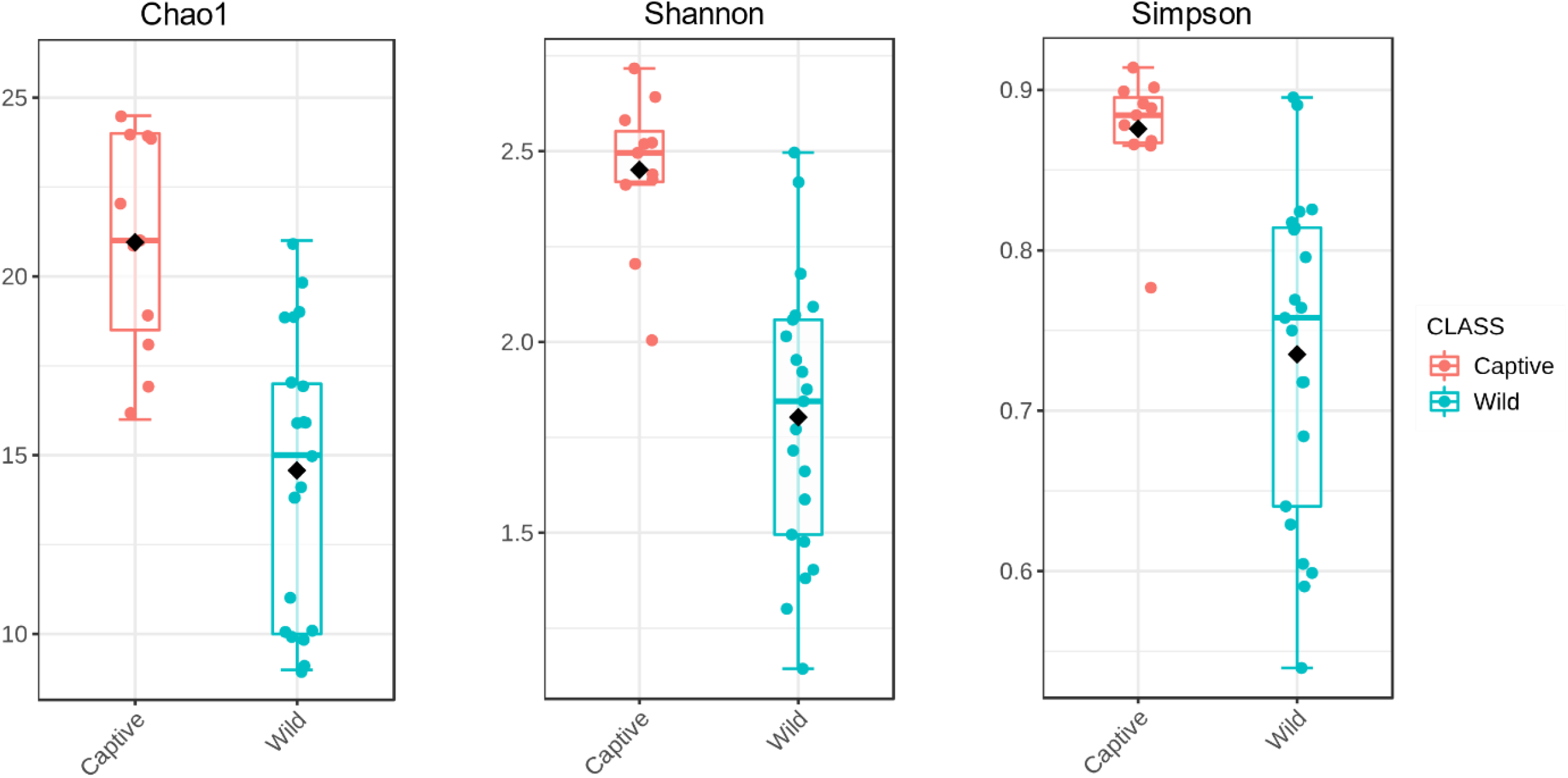
**Chao1, Shannon, and Simpson diversity** estimate differences in diversity between multiple groups. The box and whisker plots illustrate the differences in diversity between the captive and wild beetle microbiomes. Since each estimate has drawbacks in the way that diversity is calculated, these three indices were chosen to reduce individual bias in each.

**Figure 3:**
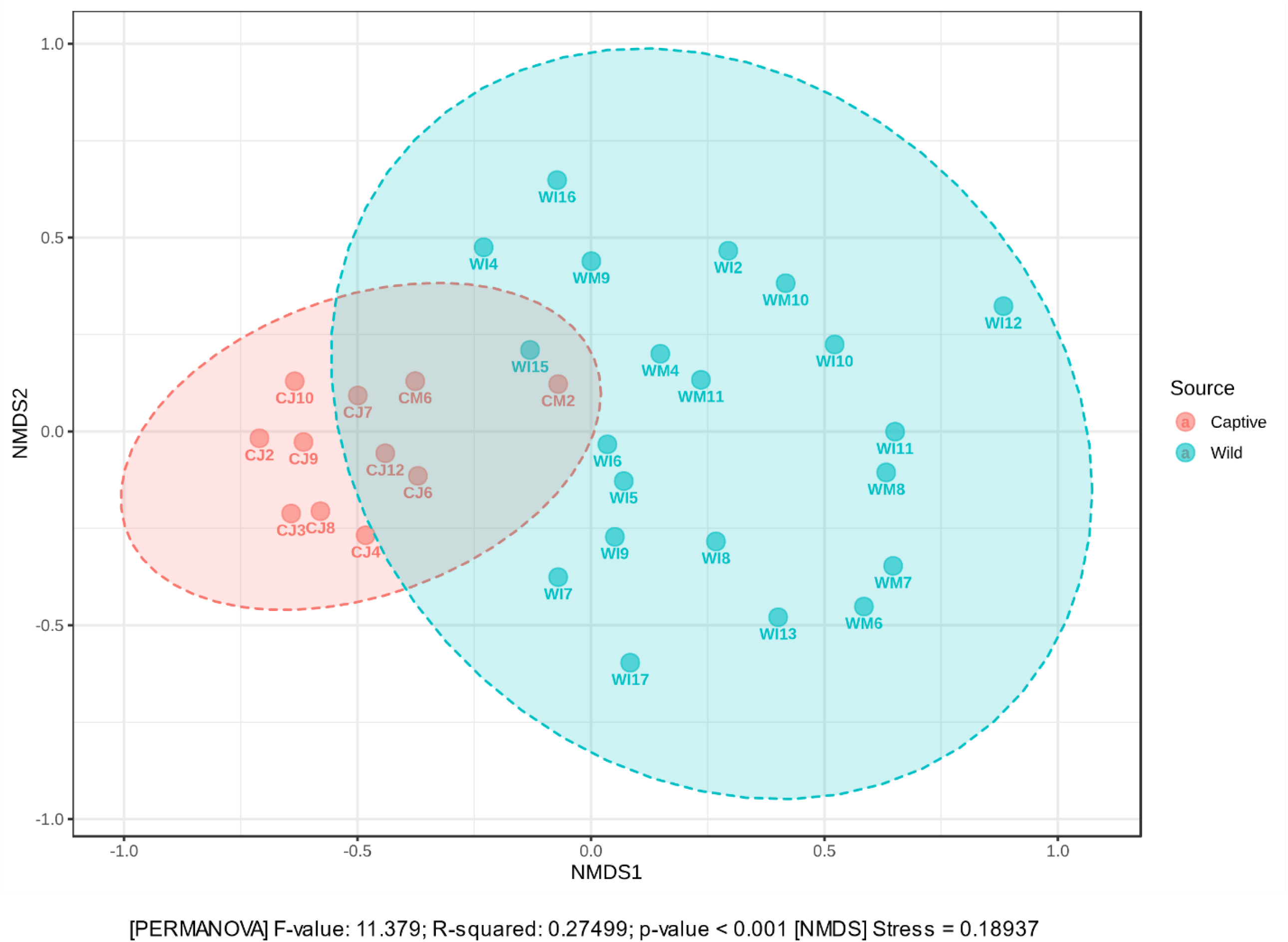
**Non-metric multidimensional scaling analysis** used the Bray-Curtis distance and PERMANOVA to rank individual beetle microbiomes among each other. Ellipses represent a 95% confidence interval around the centroid. Each beetle microbiome with over 1000 reads is represented by a point on the graph.

We used Linear Discriminant Analysis Effect Size (LEfSE) to identify 24 ASVs that were significantly different in abundance between the two groups of the 250 ASVs identified (**Figure 4**). We found that there were more ASVs overrepresented in the captive beetles (17) than in the wild beetles (7). We conducted a nBLAST search of all the ASVs the LEfSE indicated as significant. There were >98% matches of several ASVs: ASV 74 as *Sebaldella termitidis* NCTC 11300 (NCBI Taxon ID 826), ASV 19 as *Candidatus Nitrospirae nitrificans* COMA2 (NCBI Taxon ID 1742973), and ASV 62 as *Sphingorhabdus sp* GY_G (NCBI Taxon ID 2292257) (**Table 2**). ASV 4 was classified as *Staphylococcus* and had 100% match with 10 genomes belonging to *Staphylococcus scuiri*. We found some ASVs that were close relatives to several other insect microbiome components (74, 4), as well as environmental water (19), water treatment sludge samples (62, 4, 19, 72), and contaminated natural environments (4).

**Figure 4:**
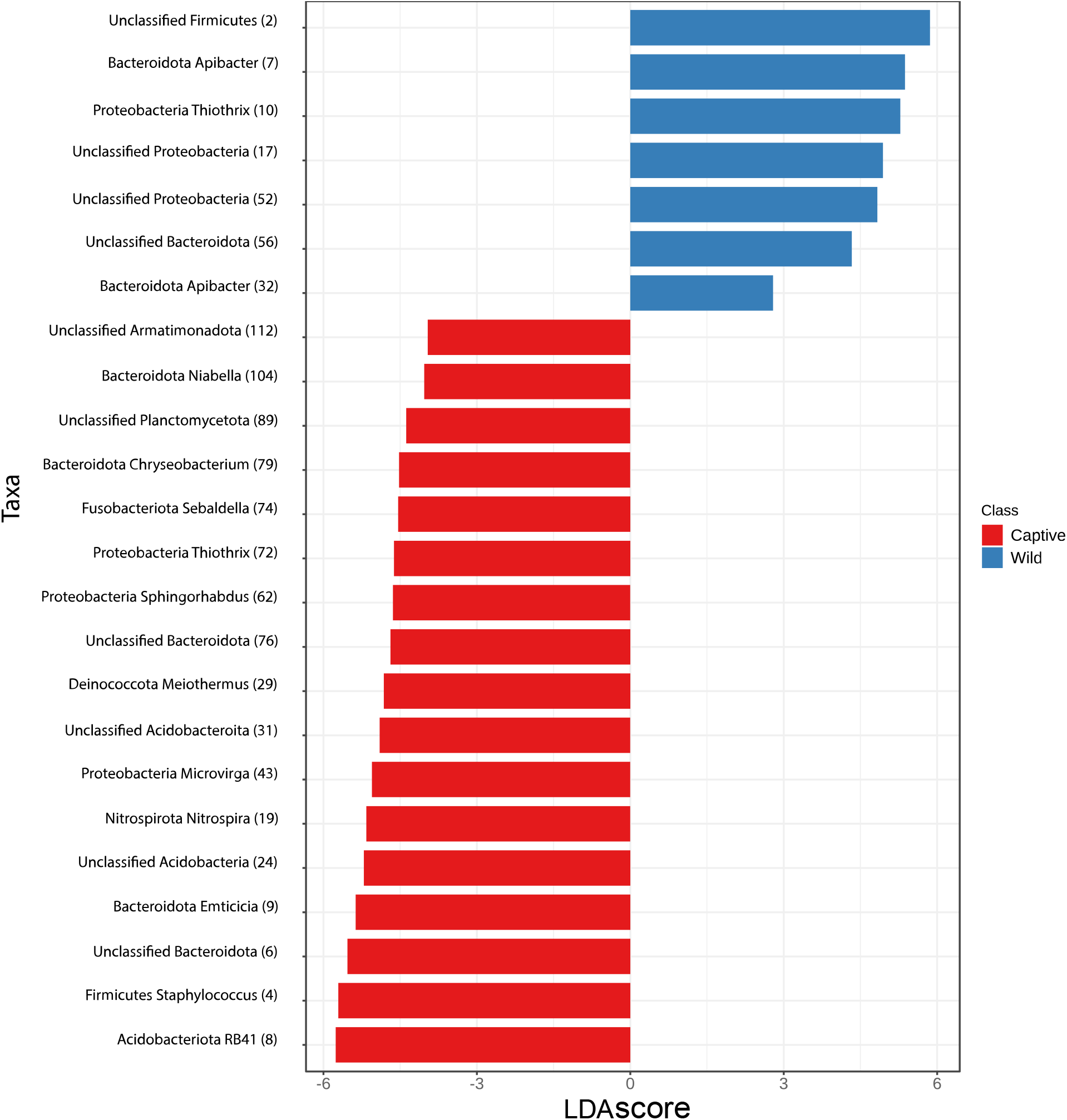
**Linear Discriminant Analysis Effect Size (LEfSE) tool as a bar plot** identifies ASVs that occur in significantly different amounts across multiple samples to explain differences between the captive and wild beetle microbiomes. Each ASV has the assigned phylum and genus, if available, followed by the ASV number to differentiate between ASVs with the same taxonomy assigned to it. Horizontal bars represent significant (P-value < 0.05) LDA scores.

**Table 1:**
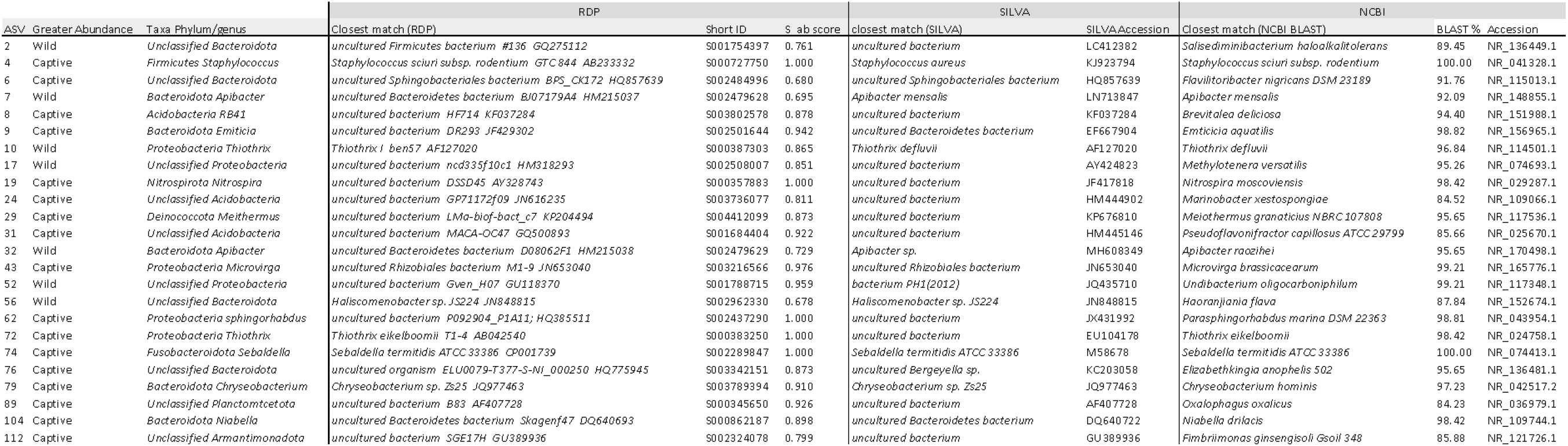
Closest matched taxa of each significant ASV. Closest matches were taken from the Ribosomal Database Project, Silva, and NCBI respectively. Significance was determined by the LefSE.

## Discussion

To the best of our knowledge, this is the first study to document the microbiome of Comal Springs riffle beetles both in the wild and in a captive setting and found noticeable differences between them. microbiomes were dominated by *Bacteroidota*, Firmicutes, and Proteobacteria phyla, as is common in insect species (Yun *et al*., 2014). We found 250 individual ASVs in the microbiomes and identified 24 ASVs that were in significantly different abundances between the groups (Figure 4).

Two phyla with large differences in abundance in the microbiomes, *Nitrospirae* and *Acidobacteria*, both have specific environmental preferences that may indicate differences in the environmental conditions between the wild and refugium habitats. *Nitrospirae* is a phylum that is known for its metabolic versatility and role in the nitrogen cycle by detoxifying ammonia and other compounds that are potentially harmful for other forms of life (Arshad *et al*., 2017). We found that beetles from the refugium contained higher relative abundances of *Nitrospirae*, which could be indicative of varying bioavailable nitrogen in the water and/or biofilms when compared to the wild environments. There might be less nitrogen in the wild for the beetles to process, other bacteria could be fulfilling this role in wild beetles, or a completely nitrogen unrelated reason.

*Acidobacteria* is a diverse phylum of acidophiles found across a wide range of ecotypes, whose functions span from carbohydrate utilization, energy metabolism, production of exopolysaccharides, to nitrogen assimilation (Kielak *et al*., 2016, Eichorst *et al*., 2018). We were not able to find exact matches of the *H. comalensis* beetle *Acidobacteria* in publicly available genomes and as the phylum is so diverse, we can only speculate as to some probable differences between the two and why captive *H. comalensis* have higher and more diverse levels of *Acidobacteria*. Nutrients and pH can drive the types and abundance of *Acidobacteria* (Kielak *et al*., 2016), but as the pH of the water habitat of wild beetles (average 7.1) is similar to that of captive beetles (average 7.2), available nutrients are likely to drive the differences. All *Acidobacteria* genomes described have contained genes for the ammonia transporter channel family and can use both organic and inorganic N, so it is likely these bacteria play a role in helping *H. comalensis* adults process this in their gut (Kielak *et al*., 2016, Eichorst *et al*., 2018). Due to widespread inclusion of genes involved in polyketide synthesis pathways in *Acidobacteria* there is speculation that some *Acidobacteria* could produce antimicrobials, but this cannot be confirmed without study of their products (Kielak *et al*., 2016). ASV 8’s closest match was from *Acidobacteria* subdivision 4, whose members have been shown to break down both chitin (Foesel *et al*., 2013, Huber *et al*., 2014) and cellulose (Huber *et al*., 2014). In general, all subdivisions degrade xylan (Kielak *et al*., 2016).

The *Dysgonomonas* genus, which is common in insect microbiomes (Tagliavia *et al*., 2014, Deguenon *et al*., 2019) was estimated to have higher relative abundance in the wild beetles than captive ones. *Dysgonomonas* is commonly found in cellulose and hemicellulose degrading bacteria within hosts, such as termites and filth-associated insects (Xinxin *et al*., 2018, Deguenon, 2019). This could indicate that cellulose, such as wood, might play a higher role in *H. comalensis* diets than previously assumed, perhaps during the process of eating biofilms off woody debris. Based on this, we would suggest captive holding containers to include more woody items for biofilm to grow on.

The captive beetles also contained *Sebaldella terminidis* (ASV 74) that had a 100% match across RDP, SILVA, and nBLAST with *Sebaldella termitidis* ATCC 33386. This species of *Sebaldella* degrades uric acid into carbon dioxide, ammonia, and acetate, and is believed to play a role in providing nitrogen to its termite hosts in low nitrogen environments (Collins & Shah, 1986). Given the presence of other bacteria involved in the nitrogen cycle, it is possible that the available nitrogen for wild beetles may not be reflected in the refugium.

The overall diversity of the captive *H. comalensis* microbiomes are higher than the wild ones. Despite the higher bacterial diversity of the captive beetle microbiomes, their microbiomes are more similar to each other in composition compared to the wild beetle microbiomes based on the nMDS (**Figure 3**). The wild beetle microbiomes individually had lower diversity, but differed more from one another according to the distance between groups in the nMDS, which could be due to increased physical distances between beetles and variation in the environment (Huston & Gibson, 2015, Ajani *et al*., 2018, Haworth *et al*., 2019). One study found that the gut of captive rhinoceroses has a higher bacterial diversity attributed to contact with humans and interaction with man-made environments (McKenzie *et al*., 2017). While the group found that many other mammals experience decreased gut microbial diversity in captivity, the researchers speculate that host traits are partially responsible for the direction of gut microbiome changes with environmental changes. Introducing bacteria to the refugium habitat could potentially contribute to the higher diversity observed in captive beetles. The SMARC alternates between the two well water sources weekly; during alternate weeks, a section of the water pipeline remains on standby with stagnant water resting in the pipeline. As the water rests for that time period in the non-operating well pipeline, biofilms can grow within the stagnant water within the pipe, which is not an uncommon place for bacteria to colonize (Kitajima *et al*., 2021). Upon a well switch it takes approximately four hours for this water to flush through the pipes. Staff have protocols to divert as much of this water as they can without being overly wasteful and to also keep water flowing in the holding containers. Additionally, the pipeline is several decades old, so the microbial community within the pipeline itself is potentially different from the aquifer community. It is possible that the microbial community of the water is changed by biofilm shedding as it travels through the pipes from the aquifer to the beetle’s captive habitat. Studies are underway in our lab group accessing the bacterial communities in the water and biofilms of both Comal Springs and the SMARC refugium.

Gray & Engel (2013) found that the microbial communities along the freshwater/saline-water interface in the karst groundwater of the Edwards (Balcones Fault Zone) Aquifer varied between different sites and well depths due to the geochemical stratification. Therefore, it is possible that the water used in the SMARC facility harbors a distinct microbial community compared to the water at the Comal Springs, which could explain some of the differences between the microbiome of wild and captive beetles. While the waters wild *H. comalensis* encounter at the spring openings of the aquifer emerge from these depths, this water might have a different microbial community and chemistry compared to the SMARC well water that is pulled from 320-402 feet in the aquifer.

Not only could the captive *H. comalensis* be exposed to additional bacteria from the water, but they could also receive exposure from human interactions, including handling of the substrates they eat or the beetles themselves. A nBLAST search of the *Staphylococcus* 16S rRNA gene against the genomes within the IMG, SILVA, RDP, and nBLAST databases returned a 100% match to several *Staphylococcus sciuri* isolates. *S. sciuri* was first described in 1976 from strains isolated from animal and human skin (Kloos *et al*., 1976). It is a common bacterium in animals and found in environmental sources such as soil and water (Dakić *et al*., 2005, Nemeghaire *et al*., 2014), though increasingly it has adapted to hospital environments (Dakić *et al*., 2005). It is hypothesized that *S. sciuri* colonized via environmental dust in the hospitals (Dakić *et al*., 2005). *S. sciuri* has also been linked as an opportunistic pathogen in compromised corals and influenced detrimental biofilm formation (Divya *et al*., 2018). It is a known opportunistic pathogen that contains a wide range of virulence and resistance genes, but is often considered commensal or harmless (Nemeghaire *et al*., 2014). More study is needed to determine what role *S. sciuri* plays for *H. comalensis*, but we urge caution as it was mainly found in captive beetles.

This study shows that there are clear differences between the microbiomes of the wild and captive *H. comalensis* namely the presence of *Nitrospirae* and *Acidobacteria* as well as some evidence for influence from environmental and human interaction factors. The data from the wild *H. comalensis* microbiomes can be used as the early stages of a baseline microbiome for adult beetles. The adult baseline microbiome can therefore help future studies guide beetle microbiomes toward a beneficial and healthy gut. Detailed work uncovering the metabolic pathways between *H. comalensis* and their microbiomes will be required to pinpoint metabolites and genes involved in normal functioning beetles.

## Acknowledgements

This work was supported by a Cooperative Agreement Award F19AC00330 with the U.S. Fish and Wildlife Service. The findings and conclusions in this article are those of the authors and do not necessarily represent the views of the U.S. Fish and Wildlife Service.

